# Clusterin secretion is attenuated by pro-inflammatory cytokines in culture models of cartilage degradation

**DOI:** 10.1101/2020.05.05.078105

**Authors:** Csaba Matta, Christopher R. Fellows, Helen Quasnichka, Adam Williams, Bernadette Jeremiasse, David Allaway, Ali Mobasheri

## Abstract

Proteomic studies have implicated clusterin as a potential biomarker of osteoarthritis (OA). However, there are two isoforms of clusterin with opposing functions, and their roles in OA have not previously been clarified. The secreted form of clusterin (sCLU) is a cytoprotective extracellular chaperone which prevents protein aggregation and enhances cell proliferation and viability, whereas nuclear clusterin (nCLU) acts as a pro-death signal. In this study, we focused on the role of sCLU and used established, pathophysiologically relevant, *in vitro* culture models to validate this potential biomarker of cartilage degradation. The secretome of equine cartilage explants, osteochondral biopsies and chondrocytes was analysed by western blotting for released sCLU, cartilage oligomeric protein (COMP) and matrix metalloproteinases (MMP) 3 and 13, following treatment with or without pro-inflammatory cytokines interleukin-1β (IL-1β) and tumour necrosis factor-α (TNF-α). The amount of sulphated glycosaminoglycans (sGAG) released into the medium was determined by dimethylmethylene blue (DMMB) analysis. Clusterin mRNA expression was quantified by real-time PCR. MMP-3, MMP-13, COMP and sGAG released from explants and osteochondral biopsies was elevated with cytokine treatment, confirming cartilage degradation in these models. Release of sCLU was attenuated with cytokine treatment in all three *in vitro* models. Expression of clusterin mRNA in cartilage explants and chondrocytes was down-regulated 7-days post cytokine stimulation. Cytokine stimulation attenuated expression and secretion of sCLU, therefore potentially limiting the cytoprotection which sCLU provides. These observations further implicate sCLU as having a role in OA, and diagnostic value as a potential biomarker for cartilage degradation.

## INTRODUCTION

Osteoarthritis (OA), the leading cause of physical disability in adults, is a degenerative disease of synovial joints characterised by inflammation, bone remodelling, and progressive degradation of articular cartilage^1^. There is a need for robust biomarkers which identify the onset of disease, enable monitoring of progression, and efficacy of potential therapeutic interventions. Identification of an early marker of OA is paramount since long periods of asymptomatic degeneration of the joint can occur, and patients often present only once damage to the cartilage is irreversible^2^. Newly synthesised molecules secreted by chondrocytes in diseased tissue, along with products of native cartilage matrix degradation, provide potential biomarkers of OA. To date, matrix degradation products have not yet yielded clinically robust biomarkers of early OA^3^. We have instead focused on secreted molecules which are uniquely expressed in early disease.

We have previously identified clusterin (CLU) as a potential biomarker of OA in an equine explant model of inflammatory joint disease^4^. CLU, also known as apolipoprotein J, is a secreted glycoprotein constitutively expressed in a broad spectrum of tissues, and has been functionally implicated in numerous physiological processes and age-related diseases, including OA^5^. CLU exists in two isoforms, a glycosylated secretory form (sCLU) and a non-glycosylated cytoplasmic form which can translocate to the nucleus under stress conditions and initiate apoptosis (nCLU)^6–8^, although certain studies did not confirm nuclear localisation of clusterin isoforms^7^. sCLU has a cytoprotective function and binds to numerous proteins preventing their aggregation, and initiating disposal by phagocytic cells^9–12^.

CLU was identified in the secretome of human healthy and OA cartilage explants as an endogenous protein released at significantly lower amounts from hip OA samples^13^. Lower levels of CLU were found to be associated with greater cartilage lesion size^14^. We are particularly interested in the clinical potential of sCLU biomarker in OA, due to its presence in serum and synovial fluid^15^. Alterations in CLU expression have previously been described in OA cartilage and synovial fluid^4; 15–23^. However, these studies have reported contrasting results, with both increases and decreases in CLU expression. We believe this is due to the existence of the two CLU isoforms, with opposing actions, which have not previously been distinguished in OA.

In this study, we have focused exclusively on the expression of sCLU in models of *in vitro* cartilage degradation through targeted biochemical analysis. We utilised cartilage explant, osteochondral biopsy and chondrocyte model systems to determine sCLU secretion with and without pro-inflammatory cytokines interleukin-1β (IL-1β) and tumour necrosis factor-α (TNF-α). We also used IL-1β and TNF-α in combination with dexamethasone, a steroidal anti-inflammatory drug commonly used to treat joint inflammation^24; 25^, to ascertain the potential of sCLU to monitor drug efficacy. Cartilage degradation in these models was determined by the quantification of known markers matrix metalloproteinases (MMP) 3 and 13, cartilage oligomeric protein (COMP) and sulphated glycosaminoglycan (sGAG)^4^. Our aim was to validate sCLU, a protein previously implicated in OA pathophysiology, as a biomarker of early OA using three *in vitro* culture model systems.

## METHODS

All reagents were supplied by Sigma-Aldrich UK unless otherwise stated.

### Cartilage and osteochondral explant culture

Macroscopically normal equine articular cartilage was obtained from the weight bearing region of the metacarpophalangeal joint of nine horses. Animals were sourced from an abattoir and euthanized for purposes other than research. The study received the full approval of the local ethics committee and was exempt from review by animal welfare authorities as abattoir tissue was used. Six-millimetre diameter explants of cartilage of equal thickness were harvested aseptically using a sterile biopsy punch (Steifel), and 10 mm osteochondral explants were isolated using an osteochondral autograft transfer system (Arthrex). Explants were placed into low glucose (1 g/L) Dulbecco’s Modified Eagle’s Medium (DMEM) supplemented with 200 mM glutamine, 500 μg/mL penicillin, 500 IU/mL streptomycin and 50 μg/mL gentamicin for 20 minutes before being washed twice in low glucose DMEM supplemented with 200 mM glutamine, 200 μg/mL of penicillin, 200 IU/mL streptomycin and 50 μg/mL gentamicin (control medium). Excess media was removed by centrifugation and explants were weighed. One osteochondral explant or ~60 mg explanted cartilage was placed into one well of a 24-well plate, per treatment, and equilibrated overnight in 1 mL of control medium at 37°C and 5% CO2. Osteochondral explants were then cultured in 1 mL fresh medium for 14 days. Explanted cartilage was cultured in 20 μL media/mg explant for 7 days.

### Chondrocyte isolation and culture

Cartilage explants were also used for the isolation of equine chondrocytes and collected as described above. Explants were diced and then digested by sequential incubations with pronase (70 U/mL for 1 hour at 37°C; Roche) and collagenase (0.2% w/v, overnight at 37°C; Gibco). Isolated passage 0 chondrocytes were seeded at high density (105,000 cells/cm^2^) and incubated in high glucose (4.5 g/L) DMEM supplemented with 200 mM glutamine, 200 μg/mL of penicillin, 200 IU/mL streptomycin, 50 μg/mL gentamicin, 1 mM sodium pyruvate, 10 μg/mL insulin, 5 μg/mL transferrin and 6.7 ng/mL selenium (Gibco) (control medium), for 7 days.

### Experimental design

Cartilage explants and chondrocytes were subjected to the following treatments; corresponding control medium in the presence or absence of 0.1, 1.0 or 10 ng/mL TNF-α (R&D Systems) and 0.1, 1.0 or 10 ng/mL IL-1β (R&D Systems). Cartilage explants were also treated with the anti-inflammatory dexamethasone (at 100 μM). Osteochondral explants were incubated in control medium in the presence or absence of 10 ng/mL TNF-α and 10 ng/mL IL-1β. Media was collected and replenished every 3.5 days, and protease inhibitor cocktail (Halt, Thermo Fisher) was added to collected media. Monolayer chondrocytes were lysed, and protein extracted, by 50 mM Tris-HCl (pH 6.8), 1% sodium dodecyl sulphate (w/v), 10% glycerol (v/v) and protease inhibitor cocktail (Halt, Thermo Fisher). Cell lysate was syringed with a 25G needle and centrifuged at 12,000×*g* for 5 min to pellet cell debris. Cell lysate supernatant and collected media were stored at −80°C prior to analysis.

### Western blotting

Cell lysate total protein was determined by using the Pierce Bicinchoninic acid (BCA) assay (Thermo Scientific), following the instructions of the manufacturer. Media and cell lysate samples were subjected to western blotting under reducing conditions. Explant media loading volumes were normalised to explant wet weight. Cell media and cell lysate loading volumes were normalised to total cell protein. Generated by pooling samples, positive control standards (STD) were used to normalise between blots. Collagen type II standard (TII STD) was generated by extraction of cartilage explants in 4 M guanidine hydrochloride and the insoluble pellet reconstituted in 0.5 M acetic acid, and further diluted with 50 mM Tris-acetate. Samples and standards were loaded onto 4–12% Tris–glycine or 10% Bis-Tris acrylamide gels (NuPAGE, Invitrogen), then transferred onto 0.45 μm nitrocellulose membrane. Membranes were blocked with 5% (w/v) bovine serum albumin (BSA) in Tris-buffered saline 0.1 % Tween-20 (v/v) (TBS-T), or Odyssey blocking buffer (Licor), for 1 h at room temperature and then incubated overnight at 4°C with the following primary antibodies diluted in 5% BSA/TBS-T or Odyssey blocking buffer with 0.2% Tween-20; anti-clusterin (Aviva Systems Biology ARP61142_P050), anti-COMP (Abcam ab128893), anti-MMP-3 (Aviva Systems Biology ARP42042_P050), anti-MMP-13 (Aviva Systems Biology ARP56350_P050), anti-collagen type II (DSHB II-II6B3), rabbit normal IgG control (Sigma-Aldrich 12-370) or mouse normal IgG control (Sigma-Aldrich 12-371). Bound primary antibodies were detected with goat anti-rabbit or goat anti-mouse antibodies conjugated with alkaline phosphatase (Cambridge Bioscience), IRDye^®^ 680RD or IRDye^®^ 800CW (Licor). Positive bands were imaged using the LI-COR Odyssey FC imaging system (LI-COR Biosciences) or by incubation in alkaline phosphatase NBT/BICP buffer (Promega), and then quantified by densitometry analysis (Image J, NIH, USA). Blots were imaged using the Licor Odyssey FC imaging system and quantification of detected bands was performed using the Image Studio™ software. Positive cell lysate bands were normalised to total protein post transfer, detected by REVERT Total Protein Stain (Licor).

### DMMB assay

sGAG content was determined by the DMMB colorimetric assay as previously described^26^ and chondroitin sulphate C, from shark cartilage, was used as a standard. Absorbance readings were taken at a wavelength of 525 nm on a Tecan SPARK 10M plate reader.

### RT-qPCR

Cartilage explants were homogenised in TRI reagent using a dismembrator (Braun Biotech). RNA was isolated from explant homogenate and chondrocyte lysate by using the Qiagen RNeasy RNA isolation kit and following the manufacturer’s instructions. First strand cDNA was then synthesized using SuperScript^®^ III reverse transcriptase (Thermo Fisher), and real-time polymerase chain reaction (RT-PCR) conducted using the Techne Prime Pro thermal cycler with GoTaq qPCR Master Mix (Promega). RT-qPCR reactions were carried out in 10 μL volumes containing 3.5 mM MgCl2, 200 μM dNTPs, 0.3 μM of sense and antisense primers, 0.025 U/μL GoTaq polymerase enzyme and 1:66,000 SYBR Green-1 (Thermo Fisher). Clusterin (CLU, NM_001081944) forward CTA-CTT-CTG-GAT-CAA-CGG-TGA-CC and reverse CGG-GTG-AAG-AAT-CTG-TCC-T primers produced a 144 bp product. Clusterin gene expression was normalised to the housekeeping gene ribosomal protein S18 (RPS18, XM_001497064) using forward CAC-AGG-AGG-CCT-ACA-CGC-CG and reverse AGG-CTA-TCT-TCC-GCC-GCC-CA primers (119 bp product).

### Statistics

Statistical tests were performed using the ‘R’ statistical package (CRAN). Data sets were checked for Gaussian distribution using Shapiro-Wilks tests and equal variance using Bartlett k^2^ tests; log transformations were conducted to correct normality or unequal variance if required. Tests for statistical significance were performed using Student’s paired *T*-test and One-way ANOVA with Tukey’s HSD pair-wise comparisons. Effect size was calculated using Cohen’s d test (with effect sizes classified as small (d ≥ 0.2), medium (d ≥ 0.5), large (d ≥ 0.8) and very large (d ≥ 1.3)^27^. Mean values are reported as ± standard error of the mean (SEM).

## RESULTS

### sGAG release from cartilage explants is elevated by cytokine stimulation

To validate our *in vitro* model of cartilage degradation, we first looked at the sulphated glycosaminoglycan (sGAG) content of the explant secretome using DMMB analysis. sGAG release was increased 3-fold following stimulation with 10 ng/mL IL-1β and TNF-α compared with the control with a very large effect size (fold change increase: FC=2.98; p=0.0058, d=2.3561; Figure 1A). Levels of the cartilage degradation marker cartilage oligomeric matrix protein (COMP) in the explant secretome were increased 2-fold with 10 ng/mL IL-1β and TNF-α stimulation *versus* the control (FC=1.94, p=0.0218, d=1.7963; Figure 1B), as revealed by western blot analysis, demonstrating the catabolic effect of cytokine treatment in this model. Uncropped gel images for COMP are shown in Supplementary Figure S1.

**Figure 1.**
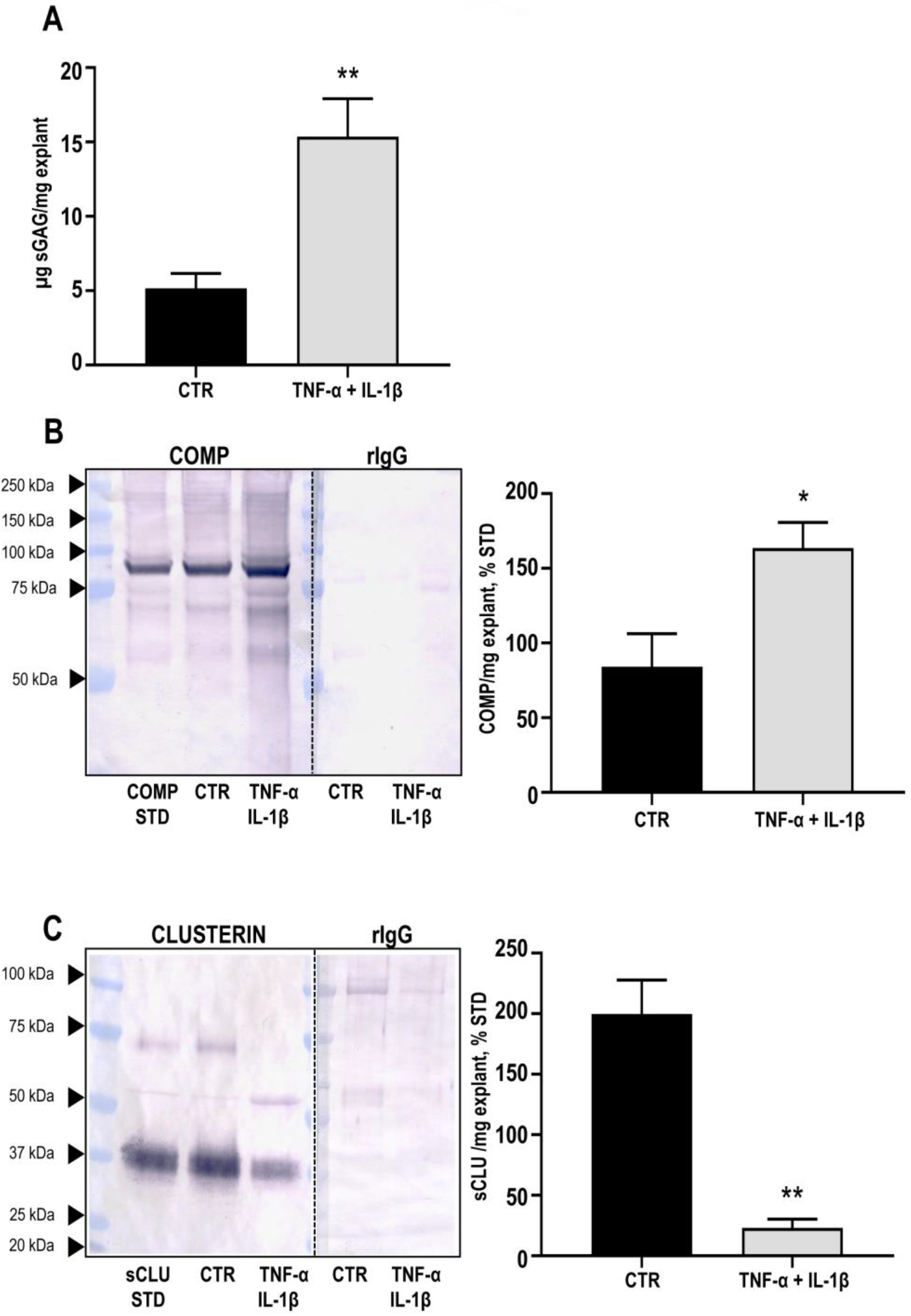
Release of sulphated glycosaminoglycans (sGAG), cartilage oligomeric matrix protein (COMP), and secreted isoform of clusterin (sCLU) into the secretome of articular cartilage explants, following stimulation with 10 ng/mL TNF-α and 10 ng/mL IL-1β for 7 days. **(A)** sGAG quantified by DMMB analysis. **(B)** COMP, **(C)** sCLU, and rabbit IgG (rIgG) controls, detected by western blotting and quantified by densitometry. Data shown are mean ± SEM, *n*=5. * and ** indicate significant differences (*p*<0.05 and *p*<0.01, respectively) as determined by Student’s *T* test versus the control (CTR).

### Clusterin release from cartilage explants is attenuated by cytokine stimulation

Mature sCLU is secreted as a heterodimeric protein with an apparent molecular weight (MW) of 75–80 kDa. Under reducing conditions, extracellular sCLU appears as α- and β-chains of 34–36 kDa and 37–39 kDa, respectively^7^. The α- and β-bands usually appear as a smear, rather than sharp bands, due to varying degrees of glycosylation^8^. Intracellular forms of clusterin include the 60 kDa precursor secretory clusterin (psCLU), the alternatively spliced 49 kDa precursor nuclear clusterin (pnCLU), and its 55 kDa post-translationally modified nuclear form (nCLU)^6; 8^.

Western blot analysis of the articular cartilage explant secretome in reducing conditions identified the glycosylated α- and β-chains of extracellular secreted sCLU between 34–39 kDa (Figure 1C). Cartilage explants stimulated for 7 days with 10 ng/mL IL-1β and TNF-α showed a 9-fold reduction in sCLU release compared to control (FC=8.69; p=0.00603, d=1.2455). Uncropped gel images for CLU are shown in Supplementary Figure S1.

### Clusterin release from osteochondral explants is attenuated by cytokine stimulation

sGAG release from ostrochondral explants was also increased 3-fold following stimulation with 10 ng/mL IL-1β and TNF-α compared with the control, with a very large effect size (FC=3.01; p=0.002, d=5.8973; Figure 2A), indicating a similar response to cartilage explants. Western blot analysis showed that sCLU expression was downregulated 363.9-fold in the osteochondral biopsy secretome following 14 days of stimulation with 10 ng/mL IL-1β and TNF-α compared to the control (p=0.0011, d=3.3291; Figure 2B). As observed in the cartilage explant secretome, the glycosylated α- and β-chains of secreted sCLU were identified in the control osteochondral secretome between 34–39 kDa. Uncropped gel images are shown in Supplementary Figure S2.

**Figure 2.**
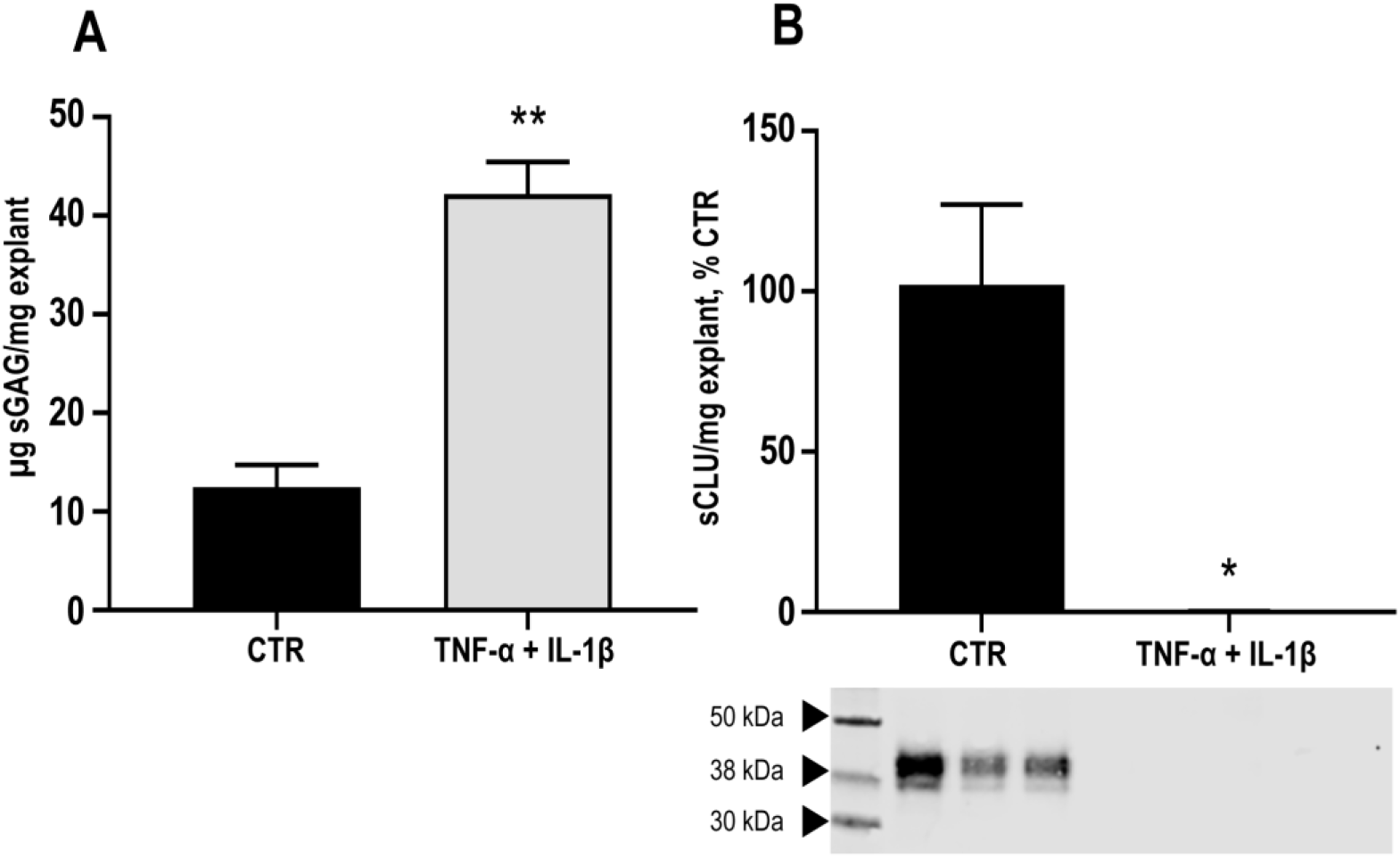
Release of sulphated glycosaminoglycans (sGAG) and secreted isoform of clusterin (sCLU) into the secretome of osteochondral explants, following stimulation with 10 ng/mL TNF-α and 10 ng/mL IL-1β for 14 days. **(A)** sGAG quantified by DMMB analysis. **(B)** sCLU, detected by western blotting and quantified by densitometry. Data shown are mean ± SEM, *n*=5. * and ** indicate significant differences (*p*<0.05 and *p*<0.01, respectively) as determined by Student’s *T* test versus the control (CTR).

### Clusterin release from cartilage explants is reduced by physiologically relevant levels of cytokine

Levels of IL-1β and TNF-α found in the synovial fluid of patients with OA are in the region of 0.1 ng/mL^28; 29^. We therefore aimed at determining whether a lower concentration of IL-1β and TNF-α had any effect on clusterin release in this model. sGAG release into the explant secretome was gradually increased by 0.1, 1.0 and 10 ng/mL IL-1β and TNF-α treatment compared with the control (fold change increases: FC=2.28, FC=3.55, FC=3.67; p=0.0283, p=0.0067, p=0.0044; and d=1.4534, d=1.746, d=1.8914, respectively; Figure 3A). In contrast, clusterin release was significantly reduced following 0.1, 1.0 and 10 ng/mL IL-1β and TNF-α treatment *versus* the control explants (fold change decreases: FC=1.32, FC=2.62, FC=6.3; p=0.0027, p=0.0158, p<0.0001; and d=1.1179, d=2.0148, d=5.9987, respectively). Uncropped gel images are shown in Supplementary Figure S3.

**Figure 3.**
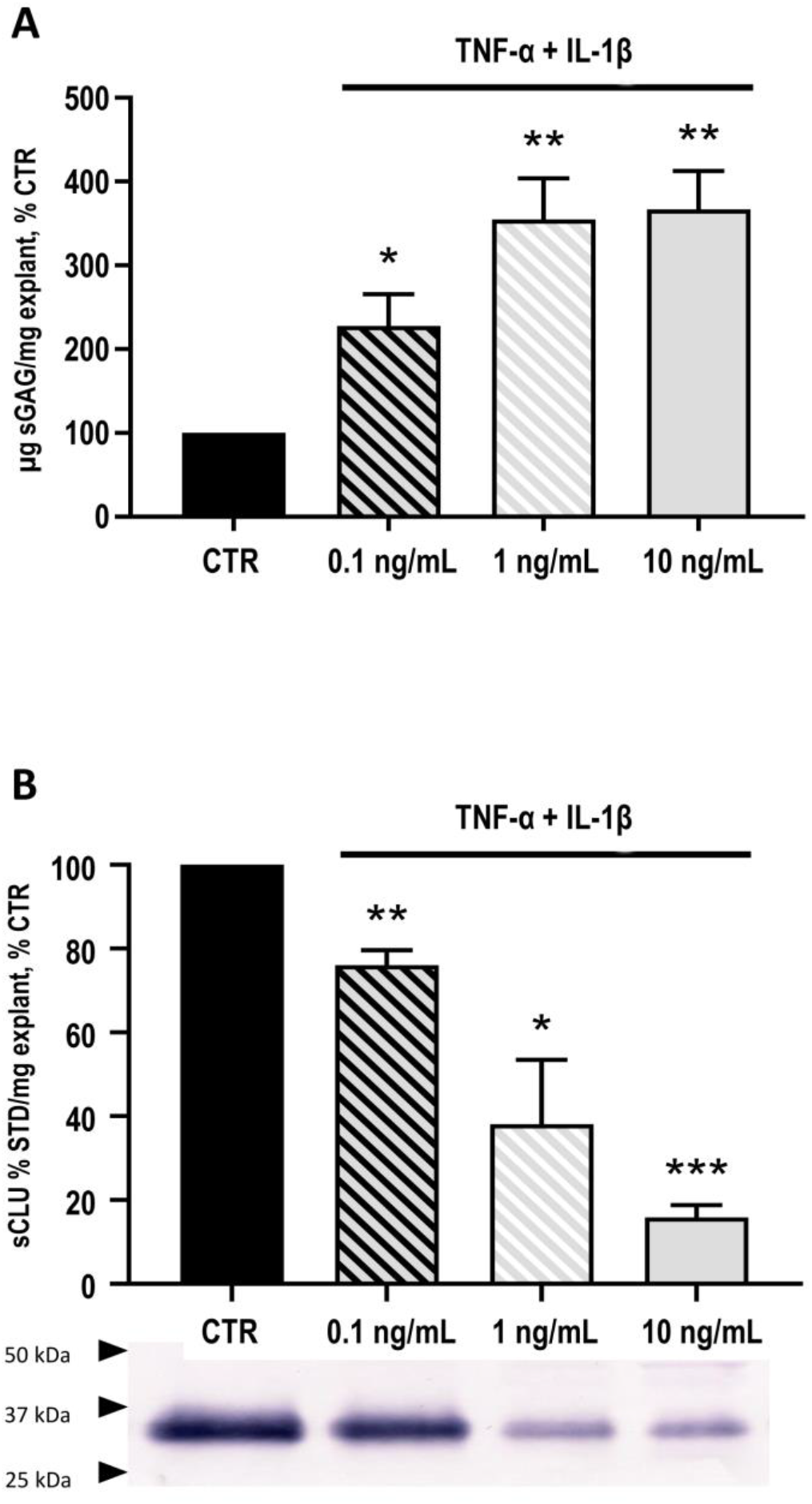
Release of sulphated glycosaminoglycans (sGAG) and secreted isoform of clusterin (sCLU) into the secretome of cartilage explants, following stimulation for 7 days with 0.1, 1 or 10 ng/mL TNF-α and 0.1, 1 or 10 ng/mL IL-1β. **(A)** sGAG quantified by DMMB analysis. **(B)** sCLU, detected by western blotting and quantified by densitometry. Data shown are mean ± SEM, *n*=3. *, ** and *** indicate significant difference (*p*<0.05. *p*<0.01 and *p*<0.001, respectively) as determined by hypothetical mean *T* test versus the control (CTR).

### Clusterin release from cartilage explants is unaffected by MMP inhibition

To determine whether the loss of clusterin release upon pro-inflammatory cytokine stimulation was due to increased proteolytic activity in the cartilage explant, and/or degradation of secreted clusterin by matrix proteases, explants were treated with 10 ng/mL TNF-α, IL-1β, and 100 μM dexamethasone in combination. Dexamethasone is an anti-inflammatory steroidal drug known to reduce MMP activity in cartilage^25^. Release of clusterin form cartilage explants, and cartilage explant clusterin gene expression were both significantly attenuated by treatment with IL-1β and TNF-α with very large effect sizes (fold change reduction: FC=109.61, p=0.0009, d=4.8357; and FC=596.57 p=0.0002, d=3.0197, versus the control, respectively; Figure 4A–B), and the same effect was observed when IL-1β and TNF-α were applied in combination with dexamethasone (fold change reduction: FC=79.44, p=0.0009, d=4.4183 and FC=204.65, p=0.0005, d=3.01, *versus* the control, respectively). In contrast, whilst IL-1β andTNF-α significantly increased MMP13 and MMP3 release into the explant secretome (FC=10.73 increase, p=0.0004, d=6.9966 and FC=4.24 increase, p=0.002, d=4.8831, *versus* the control, respectively; Figure 4C–D), co-application of dexamethasone with the cytokines abolished the increased levels of both metalloproteinases in this explant model (FC=9.61 reduction, p=0.021, d=5.59 and FC=1.95 reduction, p=0.041, d=3.37, *versus* pro-inflammatory cytokine treatment, respectively; Figure 4C–D). These data indicate that loss of secreted clusterin is unlikely due to enhanced matrix protease activity since inhibition of MMP-3 and MMP-13 by dexamethasone does not rescue cytokine-induced reduction of either clusterin release or clusterin gene expression. Uncropped gel images are shown in Supplementary Figure S4.

**Figure 4.**
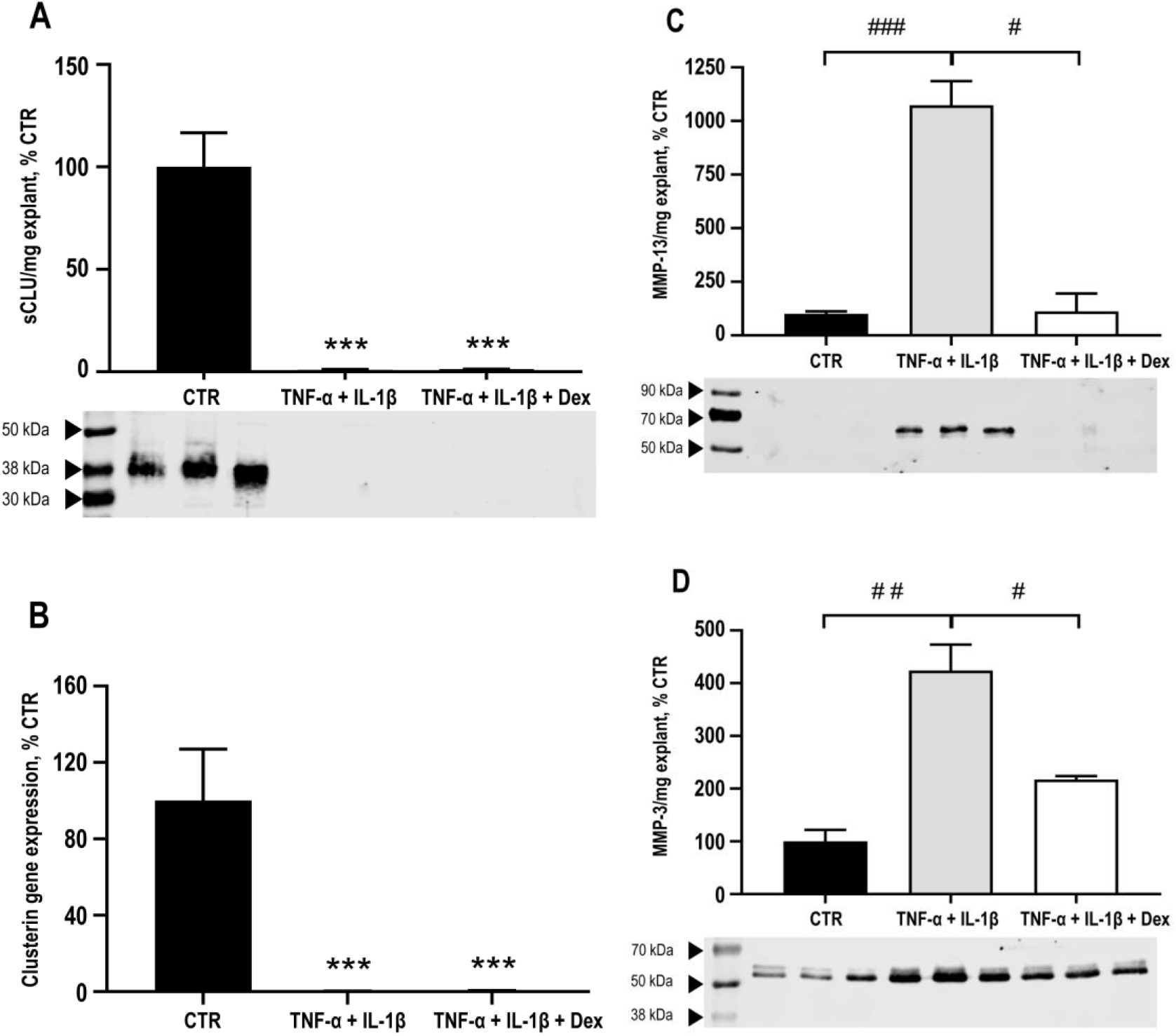
Clusterin (CLU) expression in cartilage explants, and release of secreted isoform of clusterin (sCLU) and matrix metalloproteinase (MMP) into the secretome of cartilage explants, following stimulation for 7 days with 10 ng/mL TNF-α and 10 ng/mL IL-1β ± 100 μm dexamethasone (DEX). **(A)** CLU gene expression quantified by real-time RT-qPCR analysis and normalised to ribosomal protein S18 (RPS18). **(B)** sCLU, **(C)** MMP-3 and **(D)** MMP-13 detected by western blotting and quantified by densitometry. Data shown are mean ± SEM, *n*=3. *** indicates significant difference (*p*<0.001) as determined by one-way ANOVA versus the control (CTR). #, ## and ### indicate significant difference (*p*<0.05. *p*<0.01 and *p*<0.001, respectively) as determined by one-way ANOVA versus TNF-α + IL-1β.

### Clusterin secretion from chondrocytes is reduced by cytokine stimulation

Next, we looked at clusterin release in primary un-passaged equine articular chondrocytes. Cells were cultured for 7 days in serum-free medium at high density. Secretion of sGAG and collagen type II was maintained in the control cells for the duration of the experiment, confirming the chondrocyte phenotype, as revealed by DMMB assay and western blot analysis of the chondrocyte secretome, respectively (Figure 5A–B). The α- and β-chains of sCLU were present in the chondrocyte secretome, and sCLU secretion was down-regulated by almost 4-fold with 10 ng/mL IL-1β and TNF-α stimulation, compared to the control (FC=3.79 reduction, p=0.0138, d=1.62), as revealed by western blot analyses (Figure 5C). Treatment with different concentrations (0.1, 1 and 10 mg/mL) of the cytokines resulted in a dose-dependent trend of reduced sGAG secretion (Figure 5D). A gradually attenuated release of sCLU (Figure 5E) was detected on western blots performed on the secretome following stimulation with different concentrations of the pro-inflammatory cytokines, similar to what has been observed with cartilage explants. Uncropped gel images are shown in Supplementary Figure S5.

**Figure 5.**
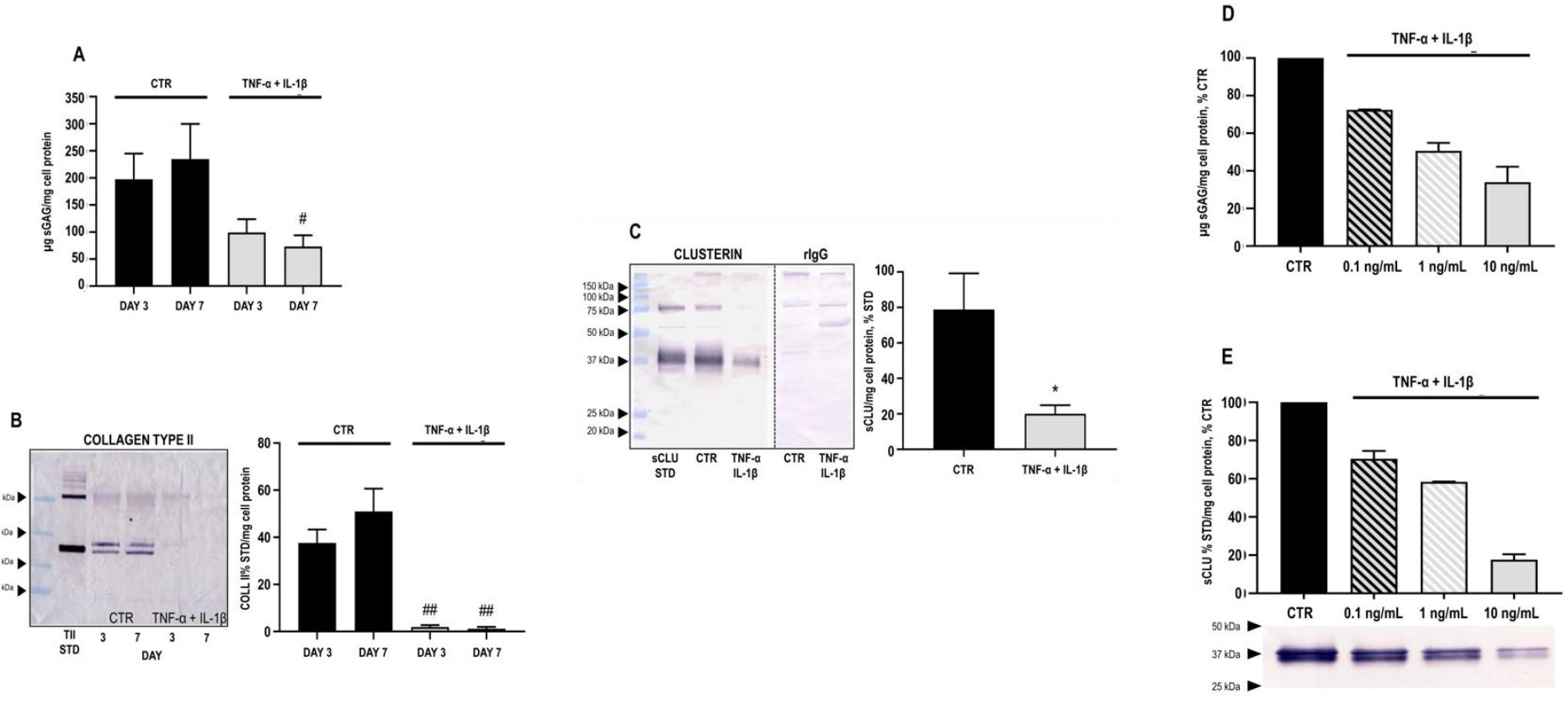
Release of sulphated glycosaminoglycans (sGAG), collagen type II, and clusterin (CLU) into the secretome of un-passaged primary articular chondrocytes, following stimulation for 3 or 7 days with TNF-α and IL-1β. **(A)** Release of sGAG following stimulation for 3 or 7 days with 10 ng/mL TNF-α and 10 ng/mL IL-1β, quantified by DMMB analysis. **(B)** Release of collagen type II (TII) following stimulation for 3 or 7 days with 10 ng/mL TNF-α and 10 ng/mL IL-1β, detected by western blotting and quantified by densitometry. Data shown are mean ± SEM, *n*=6. # and ## indicate significant difference (*p*<0.05 and *p*<0.01, respectively), as determined by one-way ANOVA versus the control (CTR) on day 3). **(C)** Release of the secreted isoform of clusterin (sCLU) into the secretome, following stimulation for 7 days with 10 ng/mL TNF-α and 10 ng/mL IL-1β, detected by western blotting and quantified by densitometry. Data shown are mean ± SEM, *n*=6. * indicates significant difference (*P*<0.05) as determined by Student’s *T* test vs the control (CTR). **(D, E)** Release of sGAG and CLU into the secretome of monolayer chondrocytes, following stimulation for 7 days with 0.1, 1 or 10 ng/mL TNF-α and 0.1, 1 or 10 ng/mL IL-1β. sGAG release was quantified by DMMB analysis; sCLU release was detected by western blotting and quantified by densitometry. Data shown are mean ± SEM, *n*=2.

### The sCLU isoform, but not the nCLU isoform, could be detected intracellularly

Western blotting performed on total lysates of primary un-passaged chondrocytes cultured for 7 days in serum-free medium at high density revealed the presence of intracellular clusterin (Figure 6A). Uncropped gel images are shown in Supplementary Figure S6. The molecular weight of the band corresponding to clusterin was confirmed at 41.1 kDa (± 0.2), indicating the presence of the intracellular CLU isoform, whilst no positive bands were detected at 49 kDa or 55 kDa (see Supplementary Figure S7), which correspond to the predicted molecular weights of nCLU. It should be noted that the anti-clusterin primary antibody employed in this study recognises an epitope in a sequence which spans both exons 6 and 7 (NM_001831.4). sCLU and nCLU both share exons 6 and 7 and differ only in the presence or absence of exons 1 and 2^7^. Treatment with the pro-inflammatory cytokines IL-1β and TNF-α at 10 ng/mL significantly reduced the levels of the intracellular CLU isoform (FC=1.45 reduction, p=0.0173, d=1.2402), compared to the control (Figure 6B).

**Figure 6.**
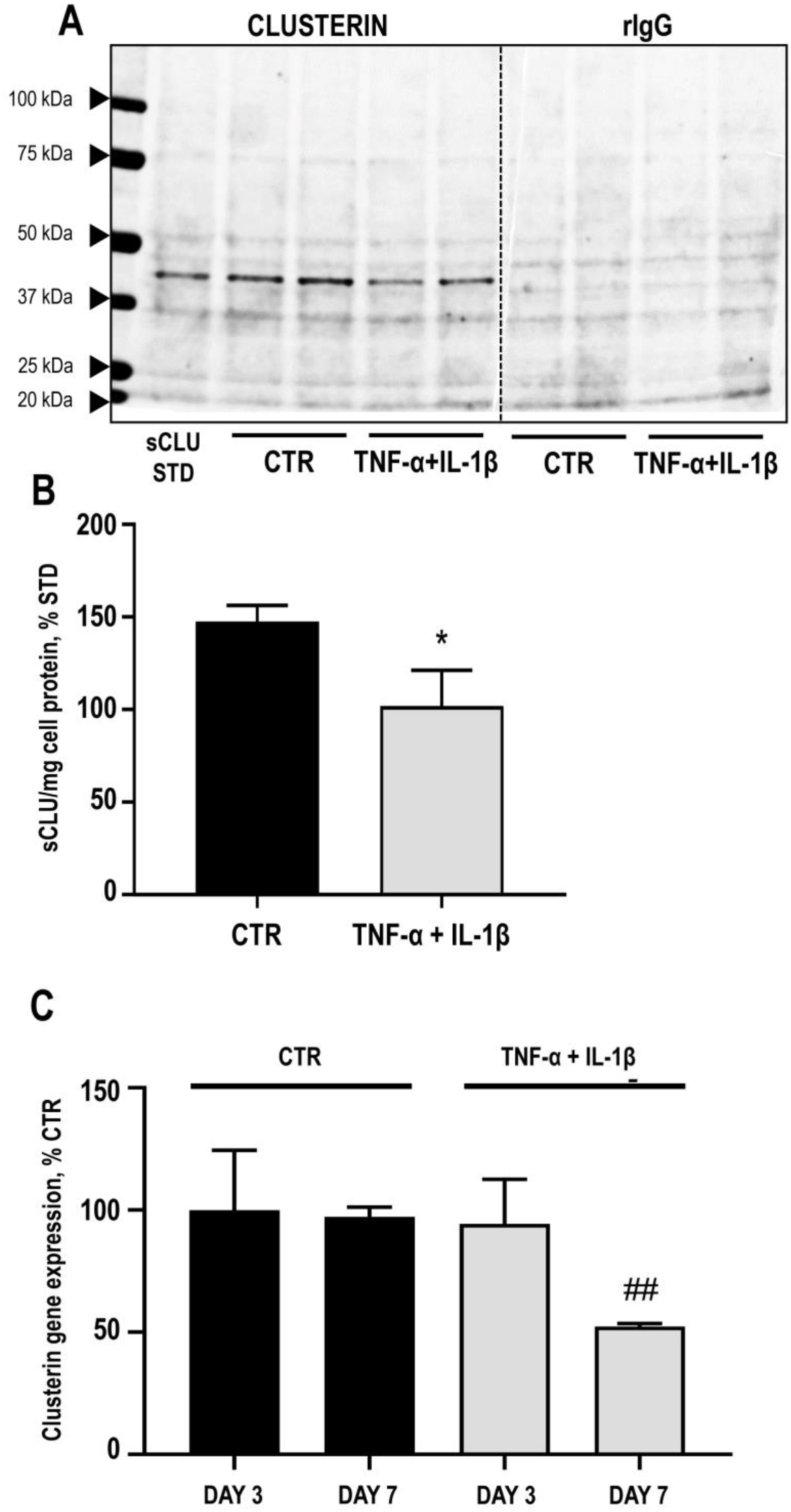
Clusterin protein and mRNA expression in un-passaged primary articular chondrocytes, following stimulation for 7 days with 10 ng/mL TNF-α and 10 ng/mL IL-1β. **(A, B)** Secreted isoform of clusterin (sCLU), and rabbit IgG (rIgG) control, detected by western blotting and quantified by densitometry; normalised to total protein loaded. Data shown are mean ± SEM, *n*=6. * indicates significant difference (*P*<0.05) as determined by Student’s *T* test versus the control (CTR). **(C)** CLU mRNA quantified by real-time RT-qPCR analysis, after being normalised to ribosomal protein S18 (RPS18). Data shown are mean ± SEM, *n*=3. ## indicates significant difference (*P*<0.01) as determined by one-way ANOVA versus the control (CTR) on day 7).

### Clusterin gene expression in chondrocyte cultures is decreased by cytokine stimulation

Clusterin gene expression was investigated using RT-qPCR analysis of primary equine articular chondrocytes cultured in a monolayer. Clusterin gene expression in chondrocyte cultures was unaffected following 3 days of stimulation with 10 ng/mL IL-1β and TNF-α; however, a 1.85-fold decrease was evident 7 days post stimulation compared to the control (p=0.005, d=8.6747), as shown in Figure 6C.

## DISCUSSION

The aim of this study was to validate secreted clusterin (sCLU) as a potential biomarker of early OA. Clusterin was identified as a candidate by our previous proteomic study, using an *in vitro* model of OA^4; 16^, and other proteomic and transcriptome studies, which analysed serum, plasma, synovial fluid and articular cartilage^13–15; 17; 18; 20–23^. Increases and decreases in clusterin levels have been reported with disease, and these contradictory findings are perhaps due to the existence of two isoforms of clusterin, and varying disease stage of donor tissue. The difference is critical due to the opposing roles of the two isoforms, and the relevance of isoform function in OA pathology. By considering only the secreted form of clusterin, and by using a targeted approach in a controlled *in vitro* model, we validated this potential marker and further elucidated its role in early articular cartilage inflammation.

We used an equine articular cartilage explant model to monitor sCLU release in response to the pro-inflammatory cytokines IL-1β and TNF-α. In order to maintain consistency with previous work from our laboratory^4; 16^, the pro-inflammatory cytokines were applied at 10 ng/mL concentration to evaluate early responses to cytokine-induced inflammation in articular cartilage. However, given that the levels of IL-1β and TNF-α found in the synovial fluid of patients with OA are in the realm of 0.1 ng/mL^28; 29^, we also aimed at determining whether lower concentrations of IL-1β and TNF-α had any effect on clusterin release in this model of cartilage degradation. The addition of the osteochondral model meant that the influence of crosstalk, at the bone–cartilage interface, on sCLU release could be ascertained^30^. Analysis of the secretome of primary chondrocytes removed the complication of pre-existing sCLU present in the cartilage matrix, enabling quantification of only newly secreted sCLU.

IL-1β and TNF-α produced by the synovial membrane are key perpetrators in the development of early OA^31^; suppressing anabolic pathways whilst stimulating catabolic enzymes, namely matrix metalloproteinases (MMPs) and aggrecanases^32; 33^. Increased levels of MMPs results in intensified proteolysis, and the progressive loss of the extracellular matrix^34^. Cytokine induction of aggrecanases leads to the loss of the large aggregating proteoglycan aggrecan, and associated release of sulphated glycosaminoglycan (sGAG) attachments^35; 36^. COMP mediates chondrocyte attachment to the ECM^37^, organises ECM assembly, and is an established marker of OA due to its correlation with disease severity and presence in synovial fluid, serum and urine^38–40^.

To validate the culture models of OA used in this study, we quantified MMP-3, MMP-13, COMP and sGAG released into the secretome of the *in vitro* cartilage explant system. Levels of MMP-3 and MMP-13 were elevated in the cartilage explant model following treatment with IL-1β and TNF-α. COMP levels also increased in the cartilage explant secretome following pro-inflammatory cytokine treatment. The sGAG content of cartilage explants was reduced with IL-1β and TNF-α stimulation confirming early-stage cartilage degradation in this inflammatory model. sGAG release into the secretome was elevated with cytokine stimulation, in both cartilage explant and osteochondral biopsy models to a similar extent, suggesting that aggrecanase-driven cartilage degradation was similarly induced in both models. The addition of the anti-inflammatory dexamethasone reduced cytokine-induced levels of MMP-3 and 13 in the cartilage explant model. These changes are in line with our previous work^4; 16^.

To quantify the newly secreted sCLU without measuring pre-existing sCLU present in the cartilage matrix, we employed monolayer cultures of freshly isolated articular chondrocytes. sGAG release into culture medium was higher in the control group, demonstrating the chondrocyte phenotype. Stimulation with IL-1β and TNF-α caused a decrease in sGAG content of the secretome, consistent with previous studies which also showed suppression of proteoglycan synthesis by cytokines^41^. Levels of both MMP-3 and MMP-13 were elevated with cytokine treatment in the secretome of the chondrocyte monolayer model.

All three culture models exhibited a marked reduction in sCLU release with IL-1β and TNF-α stimulation. We also showed that in monolayer chondrocytes and cartilage explants, clusterin gene expression was downregulated following 7-days-long stimulation with IL-1β and TNF-α, which is in line with our previous data showing a reduction in CLU precursor release^16^. The addition of dexamethasone did not affect sCLU release from cartilage explants, which suggests that the loss of clusterin is MMP-3 and 13 independent. In addition, sCLU secretion was just as marked in the osteochondral model despite less MMP release, suggesting bone-cartilage cross-talk, and that MMP levels have little influence on the fate of sCLU. Taken together, it is likely that the loss in sCLU secretion from cartilage is due to an interruption at transcriptional level, rather than degradation of extracellular sCLU. It has been reported that transforming growth factor beta (TGF-β) regulates the expression of CLU through the transcription factor c-Fos^42^. c-Fos represses CLU expression of in the absence of TGF-β, and conversely TGF-β inhibits c-Fos, resulting in increased CLU expression^42^. IL-1β and TNF-α have both been shown to exert a suppressive effect on TGF-β signalling^43^ possibly accounting for the loss of sCLU observed in our study.

Secreted clusterin may act as a cytoprotective chaperone and aids protein refolding, preventing aggregation under stress conditions, enhances cell proliferation and cell-viability, and is constitutively secreted by mammalian cells^7^. In this study, IL-1β and TNF-α interrupt expression and secretion of sCLU, and therefore the protection it offers to cells. The balance between IL-1β, TNF-α and TGF-β signalling pathways is crucial for maintenance of articular cartilage homeostasis and its disruption likely plays a key role in the pathogenesis of OA. This is perhaps, in part, due to the loss of the cytoprotective isoform of clusterin.

Both increases and decreases in clusterin levels have been reported in cartilage explants, synovium, synovial fluid and plasma of OA patients, depending on the experimental design, and varying disease stage of donor tissue^13–15; 17; 19; 20; 23^. Clusterin is increased at the transcript level in early OA cartilage, and decreased in advanced^17^. Although detectable, clusterin is significantly reduced in OA samples compared to healthy^18; 21^. Clusterin is increased in plasma, and synovial fluid, from OA patients, and peptide levels of clusterin have been suggested to be as predictive of OA progression as age^23^; and may represent activation of a compensatory, but ultimately ineffective, protective pathway^15^.

The *in vitro* explant model mimics the onset of disease, mechanisms of which may precede events observed in early clinical OA samples. Our group has previously reported decreased release of clusterin from cartilage explants stimulated with IL-1β, using high-throughput mass spectrometry, supporting findings in our current study^4; 16^. Another study also showed decreased clusterin release with IL-1β treatment in mouse femoral head cartilage^44^. Conversely, increased clusterin release was observed with IL-1β and TNF-α from bovine articular cartilage^45^.

Only in our proteomic study has the clusterin isoform been specified (15), otherwise, isoforms have not been distinguished. It is possible that the difference between the actions of sCLU and nCLU explain the contradictory picture, and increases in nCLU with simultaneous decreases in sCLU may account for the variability in clusterin levels reported. The nuclear isoform has an antithetical function to sCLU, and acts as a pro-death signal, inhibiting cell growth and survival^8; 46–48^. Cytoplasmic pnCLU is post-translationally modified in response to cell damage and cytotoxic events, and nCLU translocates to the nucleus where it induces apoptosis^6; 8^. Chondrocyte apoptosis is a key process in OA and is associated with disease progression, with late-stage exhibiting higher levels of cell death^49; 50^. Elevated levels of clusterin in late stage disease^15; 17; 19; 20; 23^ could be due to the release of intracellular nCLU following programmed cell death, whereas loss of the protective sCLU could reflect disease onset. It is essential that future studies distinguish between the two CLU isoforms.

## CONCLUSION

This study has, for the first time, identified specifically the secreted isoform of clusterin in three *in vitro* model systems of early OA. All three models consistently showed a loss of sCLU secretion with cytokine stimulation. Loss of protective sCLU in early OA could lead to increased cellular stress, which may contribute to OA progression. We propose that the secreted form of clusterin is a potential biomarker for the onset of OA.

## Supporting information

Supplemental images

## ACKNOWLEDGEMENTS

CM was supported by the European Commission through a Marie Skłodowska-Curie Intra-European Fellowship for career development (project number: 625746; acronym: CHONDRION; FP7-PEOPLE-2013-IEF). CM is supported by the Premium Postdoctoral Research Fellowship by the Hungarian Academy of Sciences, and a Bridging Fund from the Faculty of Medicine, University of Debrecen. The research was co-financed by the Thematic Excellence Programme of the Ministry for Innovation and Technology in Hungary, within the framework of the Space Sciences thematic programme of the University of Debrecen (ED 18-1-2019-0028). The research underpinning some of the work presented has received funding from a number of sources including The European Commission Framework 7 programme (EU FP7; HEALTH.2012.2.4.5-2, project number 305815; Novel Diagnostics and Biomarkers for Early Identification of Chronic Inflammatory Joint Diseases, D-BOARD) and the Innovative Medicines Initiative Joint Undertaking under grant agreement No. 115770, resources of which are composed of financial contribution from the European Union’s Seventh Framework programme (FP7/2007–2013) and EFPIA companies’ in-kind contribution. AM is a member of the Arthritis Research UK Centre for Sport, Exercise, and Osteoarthritis, funded by Arthritis Research UK (Grant Reference Number: 20194). AM is a member of the Applied Public-Private Research enabling OsteoArthritis Clinical Headway (APPROACH) consortium, a 5-year project funded by the European Commission’s Innovative Medicines Initiative (IMI). AM has received financial support from the European Structural and Social Funds through the Research Council of Lithuania (Lietuvos Mokslo Taryba) according to the activity “Improvement of researchers” qualification by implementing world-class R&D projects” of Measure No. 09.3.3-LMT-K-712 (grant application code: 09.3.3-LMT-K-712-01-0157, agreement No. DOTSUT-215). AM has also received financial support from the European Structural and Social Funds through the Research Council of Lithuania (Lietuvos Mokslo Taryba) according to the Programme “Attracting Foreign Researchers for Research Implementation,” Grant No. 0.2.2-LMT-K-718-02-0022.

## REFERENCES

1. Hunter DJ, Bierma-Zeinstra S. 2019. Osteoarthritis. Lancet 393:1745–1759.

2. Henrotin Y, Sanchez C, Bay-Jensen AC, et al. 2016. Osteoarthritis biomarkers derived from cartilage extracellular matrix: Current status and future perspectives. Ann Phys Rehabil Med 59:145–148.

3. Mobasheri A, Bay-Jensen AC, van Spil WE, et al. 2017. Osteoarthritis Year in Review 2016: biomarkers (biochemical markers). Osteoarthritis and cartilage 25:199–208.

4. Williams A, Smith JR, Allaway D, et al. 2013. Carprofen inhibits the release of matrix metalloproteinases 1, 3, and 13 in the secretome of an explant model of articular cartilage stimulated with interleukin 1beta. Arthritis research & therapy 15:R223.

5. Trougakos IP. 2013. The molecular chaperone apolipoprotein J/clusterin as a sensor of oxidative stress: implications in therapeutic approaches - a mini-review. Gerontology 59:514–523.

6. Jones SE, Jomary C. 2002. Clusterin. Int J Biochem Cell Biol 34:427–431.

7. Prochnow H, Gollan R, Rohne P, et al. 2013. Non-secreted clusterin isoforms are translated in rare amounts from distinct human mRNA variants and do not affect Bax-mediated apoptosis or the NF-kappaB signaling pathway. PLoS One 8:e75303.

8. Shannan B, Seifert M, Leskov K, et al. 2005. Challenge and promise: roles for clusterin in pathogenesis, progression and therapy of cancer. Cell Death Differ 13:12–19.

9. Baiersdorfer M, Schwarz M, Seehafer K, et al. 2010. Toll-like receptor 3 mediates expression of clusterin/apolipoprotein J in vascular smooth muscle cells stimulated with RNA released from necrotic cells. Experimental cell research 316:3489–3500.

10. Bartl MM, Luckenbach T, Bergner O, et al. 2001. Multiple receptors mediate apoJ-dependent clearance of cellular debris into nonprofessional phagocytes. Experimental cell research 271:130–141.

11. Carver JA, Rekas A, Thorn DC, et al. 2003. Small heat-shock proteins and clusterin: intra- and extracellular molecular chaperones with a common mechanism of action and function? IUBMB life 55:661–668.

12. Humphreys DT, Carver JA, Easterbrook-Smith SB, et al. 1999. Clusterin has chaperone-like activity similar to that of small heat shock proteins. The Journal of biological chemistry 274:6875–6881.

13. Fernandez-Puente P, Gonzalez-Rodriguez L, Calamia V, et al. 2019. Analysis of Endogenous Peptides Released from Osteoarthritic Cartilage Unravels Novel Pathogenic Markers. Mol Cell Proteomics 18:2018–2028.

14. Kiapour AM, Sieker JT, Proffen BL, et al. 2019. Synovial fluid proteome changes in ACL injury-induced posttraumatic osteoarthritis: Proteomics analysis of porcine knee synovial fluid. PLoS One 14:e0212662.

15. Ritter SY, Subbaiah R, Bebek G, et al. 2013. Proteomic analysis of synovial fluid from the osteoarthritic knee: comparison with transcriptome analyses of joint tissues. Arthritis and rheumatism 65:981–992.

16. Clutterbuck AL, Smith JR, Allaway D, et al. 2011. High throughput proteomic analysis of the secretome in an explant model of articular cartilage inflammation. Journal of proteomics 74:704–715.

17. Connor JR, Kumar S, Sathe G, et al. 2001. Clusterin expression in adult human normal and osteoarthritic articular cartilage. Osteoarthritis and cartilage 9:727–737.

18. Daghestani HN, Kraus VB. 2015. Inflammatory biomarkers in osteoarthritis. Osteoarthritis and cartilage 23:1890–1896.

19. Devauchelle V, Essabbani A, De Pinieux G, et al. 2006. Characterization and functional consequences of underexpression of clusterin in rheumatoid arthritis. Journal of immunology (Baltimore, Md: 1950) 177:6471–6479.

20. Fandridis E, Apergis G, Korres DS, et al. 2011. Increased expression levels of apolipoprotein J/clusterin during primary osteoarthritis. In vivo (Athens, Greece) 25:745–749.

21. Hsueh MF, Khabut A, Kjellstrom S, et al. 2016. Elucidating the Molecular Composition of Cartilage by Proteomics. J Proteome Res 15:374–388.

22. Muller C, Khabut A, Dudhia J, et al. 2014. Quantitative proteomics at different depths in human articular cartilage reveals unique patterns of protein distribution. Matrix biology: journal of the International Society for Matrix Biology 40:34–45.

23. Ritter SY, Collins J, Krastins B, et al. 2014. Mass spectrometry assays of plasma biomarkers to predict radiographic progression of knee osteoarthritis. Arthritis research & therapy 16:456.

24. Grecomoro G, Piccione F, Letizia G. 1992. Therapeutic synergism between hyaluronic acid and dexamethasone in the intra-articular treatment of osteoarthritis of the knee: a preliminary open study. Current medical research and opinion 13:49–55.

25. Huebner KD, Shrive NG, Frank CB. 2014. Dexamethasone inhibits inflammation and cartilage damage in a new model of post-traumatic osteoarthritis. Journal of orthopaedic research: official publication of the Orthopaedic Research Society 32:566–572.

26. Farndale RW, Buttle DJ, Barrett AJ. 1986. Improved quantitation and discrimination of sulphated glycosaminoglycans by use of dimethylmethylene blue. Biochimica et biophysica acta 883:173–177.

27. Sullivan GM, Feinn R. 2012. Using Effect Size-or Why the P Value Is Not Enough. J Grad Med Educ 4:279–282.

28. Manicourt DH, Poilvache P, Van Egeren A, et al. 2000. Synovial fluid levels of tumor necrosis factor alpha and oncostatin M correlate with levels of markers of the degradation of crosslinked collagen and cartilage aggrecan in rheumatoid arthritis but not in osteoarthritis. Arthritis and rheumatism 43:281–288.

29. McNulty AL, Rothfusz NE, Leddy HA, et al. 2013. Synovial fluid concentrations and relative potency of interleukin-1 alpha and beta in cartilage and meniscus degradation. Journal of orthopaedic research: official publication of the Orthopaedic Research Society 31:1039–1045.

30. Lajeunesse D, Reboul P. 2003. Subchondral bone in osteoarthritis: a biologic link with articular cartilage leading to abnormal remodeling. Curr Opin Rheumatol 15:628–633.

31. Smith MD, Triantafillou S, Parker A, et al. 1997. Synovial membrane inflammation and cytokine production in patients with early osteoarthritis. J Rheumatol 24:365–371.

32. Goldring MB, Birkhead J, Sandell LJ, et al. 1988. Interleukin 1 suppresses expression of cartilage-specific types II and IX collagens and increases types I and III collagens in human chondrocytes. J Clin Invest 82:2026–2037.

33. Goldring MB, Otero M, Tsuchimochi K, et al. 2008. Defining the roles of inflammatory and anabolic cytokines in cartilage metabolism. Ann Rheum Dis 67 Suppl 3:iii75–82.

34. Vincenti MP, Brinckerhoff CE. 2002. Transcriptional regulation of collagenase (MMP-1, MMP-13) genes in arthritis: integration of complex signaling pathways for the recruitment of gene-specific transcription factors. Arthritis research 4:157–164.

35. Huang K, Wu LD. 2008. Aggrecanase and aggrecan degradation in osteoarthritis: a review. J Int Med Res 36:1149–1160.

36. Johnson CI, Argyle DJ, Clements DN. 2016. In vitro models for the study of osteoarthritis. Vet J 209:40–49.

37. DiCesare PE, Morgelin M, Mann K, et al. 1994. Cartilage oligomeric matrix protein and thrombospondin 1. Purification from articular cartilage, electron microscopic structure, and chondrocyte binding. Eur J Biochem 223:927–937.

38. Clark AG, Jordan JM, Vilim V, et al. 1999. Serum cartilage oligomeric matrix protein reflects osteoarthritis presence and severity: the Johnston County Osteoarthritis Project. Arthritis and rheumatism 42:2356–2364.

39. Lohmander LS, Saxne T, Heinegard DK. 1994. Release of cartilage oligomeric matrix protein (COMP) into joint fluid after knee injury and in osteoarthritis. Ann Rheum Dis 53:8–13.

40. Misumi K, Tagami M, Kamimura T, et al. 2006. Urine cartilage oligomeric matrix protein (COMP) measurement is useful in discriminating the osteoarthritic Thoroughbreds. Osteoarthritis and cartilage 14:1174–1180.

41. Beekhuizen M, Bastiaansen-Jenniskens YM, Koevoet W, et al. 2011. Osteoarthritic synovial tissue inhibition of proteoglycan production in human osteoarthritic knee cartilage: establishment and characterization of a long-term cartilage-synovium coculture. Arthritis and rheumatism 63:1918–1927.

42. Jin G, Howe PH. 1999. Transforming growth factor beta regulates clusterin gene expression via modulation of transcription factor c-Fos. Eur J Biochem 263:534–542.

43. Roman-Blas JA, Stokes DG, Jimenez SA. 2007. Modulation of TGF-beta signaling by proinflammatory cytokines in articular chondrocytes. Osteoarthritis and cartilage 15:1367–1377.

44. Wilson R, Belluoccio D, Little CB, et al. 2008. Proteomic characterization of mouse cartilage degradation in vitro. Arthritis and rheumatism 58:3120–3131.

45. Stevens AL, Wishnok JS, Chai DH, et al. 2008. A sodium dodecyl sulfate-polyacrylamide gel electrophoresis-liquid chromatography tandem mass spectrometry analysis of bovine cartilage tissue response to mechanical compression injury and the inflammatory cytokines tumor necrosis factor alpha and interleukin-1beta. Arthritis and rheumatism 58:489–500.

46. Bettuzzi S, Rizzi F. 2009. Chapter 5: Nuclear CLU (nCLU) and the fate of the cell. Advances in cancer research 104:59–88.

47. Leskov KS, Araki S, Lavik JP, et al. 2011. CRM1 protein-mediated regulation of nuclear clusterin (nCLU), an ionizing radiation-stimulated, Bax-dependent pro-death factor. The Journal of biological chemistry 286:40083–40090.

48. Leskov KS, Klokov DY, Li J, et al. 2003. Synthesis and functional analyses of nuclear clusterin, a cell death protein. The Journal of biological chemistry 278:11590–11600.

49. Hwang HS, Kim HA. 2015. Chondrocyte Apoptosis in the Pathogenesis of Osteoarthritis. Int J Mol Sci 16:26035–26054.

50. Zamli Z, Adams MA, Tarlton JF, et al. 2013. Increased chondrocyte apoptosis is associated with progression of osteoarthritis in spontaneous Guinea pig models of the disease. Int J Mol Sci 14:17729–17743.

