## Supplemental images for "Clusterin secretion is attenuated by pro-inflammatory cytokines in culture models of cartilage degradation"

**This PDF file includes:**

- **Figure S1.** Original uncropped gel images of cartilage oligomeric matrix protein (COMP) and clusterin (CLU) western blots performed on cartilage explant secretome.
- **Figure S2.** Original uncropped gel images of clusterin (CLU) western blots performed on osteochondral explant secretome.
- **Figure S3.** Original uncropped gel images of clusterin (CLU) western blots performed on cartilage explant secretome after exposure to physiological levels of cytokines.
- **Figure S4.** Original uncropped gel images of clusterin (CLU) and matrix metalloprotease-3 (MMP-3) and MMP-13 western blots performed on cartilage explant secretome.
- **Figure S5.** Original uncropped gel images of collagen type II and clusterin (CLU) western blots performed on primary equine articular chondrocyte secretome after exposure to physiological levels of cytokines
- **Figure S6.** Original uncropped gel images of clusterin (CLU) western blots performed on primary equine articular chondrocyte secretome, normalised to total protein loaded
- **Figure S7.** Densitometry of intracellular isoform of clusterin (CLU) western blots performed on primary equine articular chondrocyte secretome

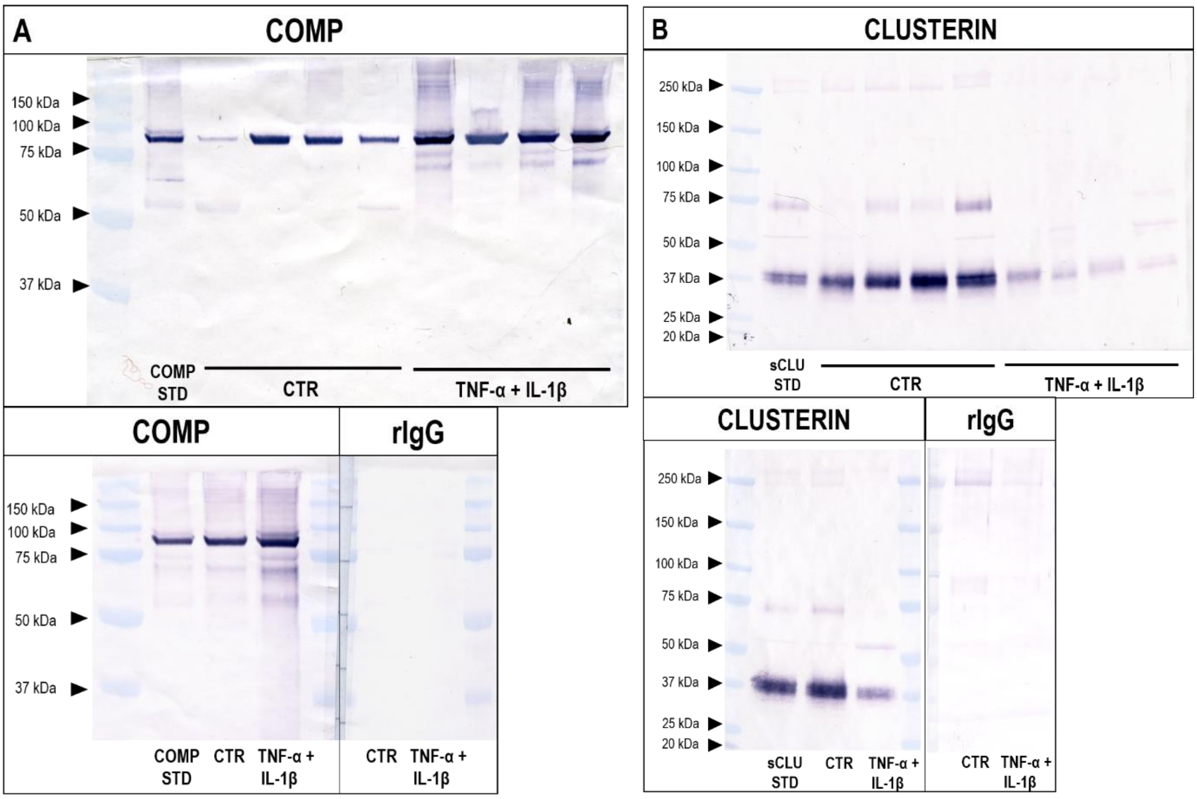


**Figure S1.** (**A**) Release of cartilage oligomeric matrix protein (COMP), and (**B**) secreted isoform of clusterin (sCLU) into the secretome of articular cartilage explants, following stimulation with 10 ng/mL TNF-α and 10 ng/mL IL-1β for 7 days. Uncropped membrane images are shown, along with rabbit IgG (rIgG) controls, detected by western blotting and quantified by densitometry.

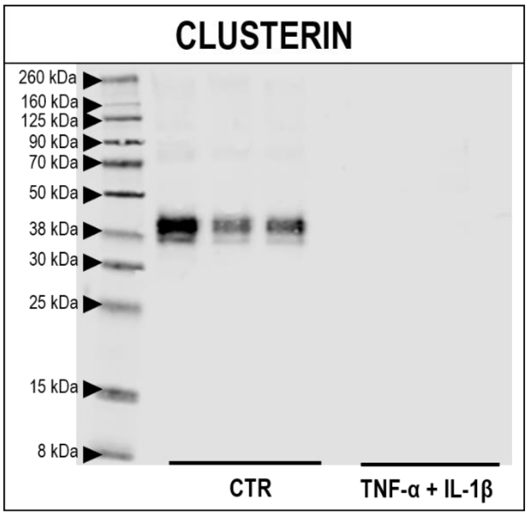


**Figure S2.** Release of secreted isoform of clusterin (sCLU) into the secretome of osteochondral explants, following stimulation with 10 ng/mL TNF-α and 10 ng/mL IL-1β for 7 days. Uncropped membrane image is shown, detected by western blotting and quantified by densitometry.

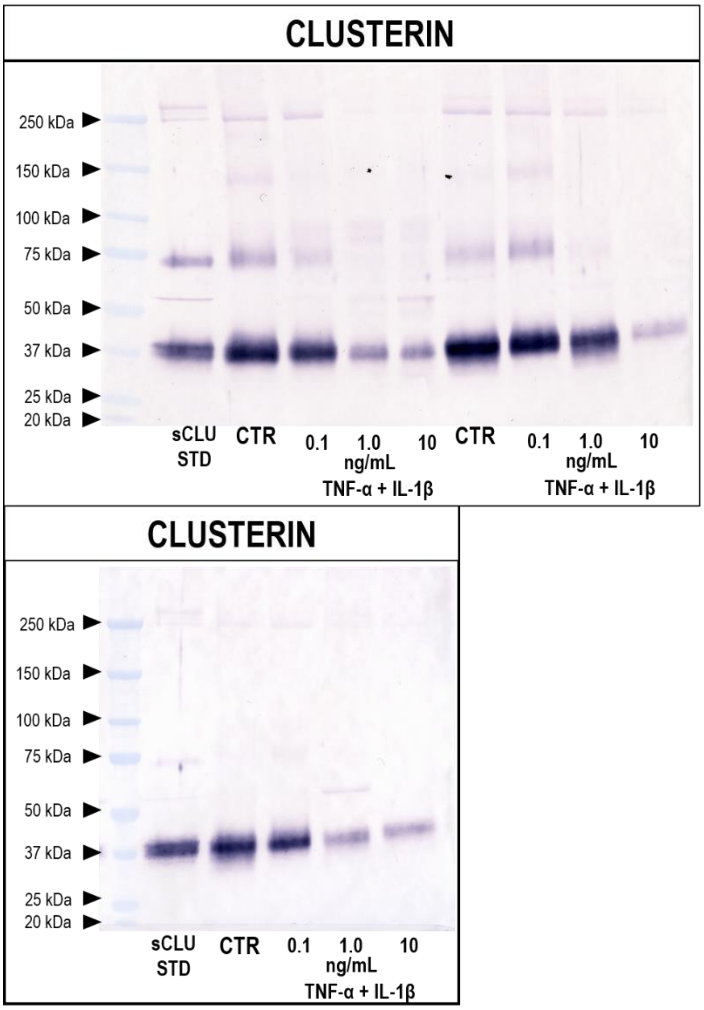


**Figure S3.** Release of secreted isoform of clusterin (sCLU) into the secretome of cartilage explants, following stimulation for 7 days with 0.1, 1 or 10 ng/mL TNF-α and 0.1, 1 or 10 ng/mL IL-1β. Uncropped membrane image is shown, detected by western blotting and quantified by densitometry.

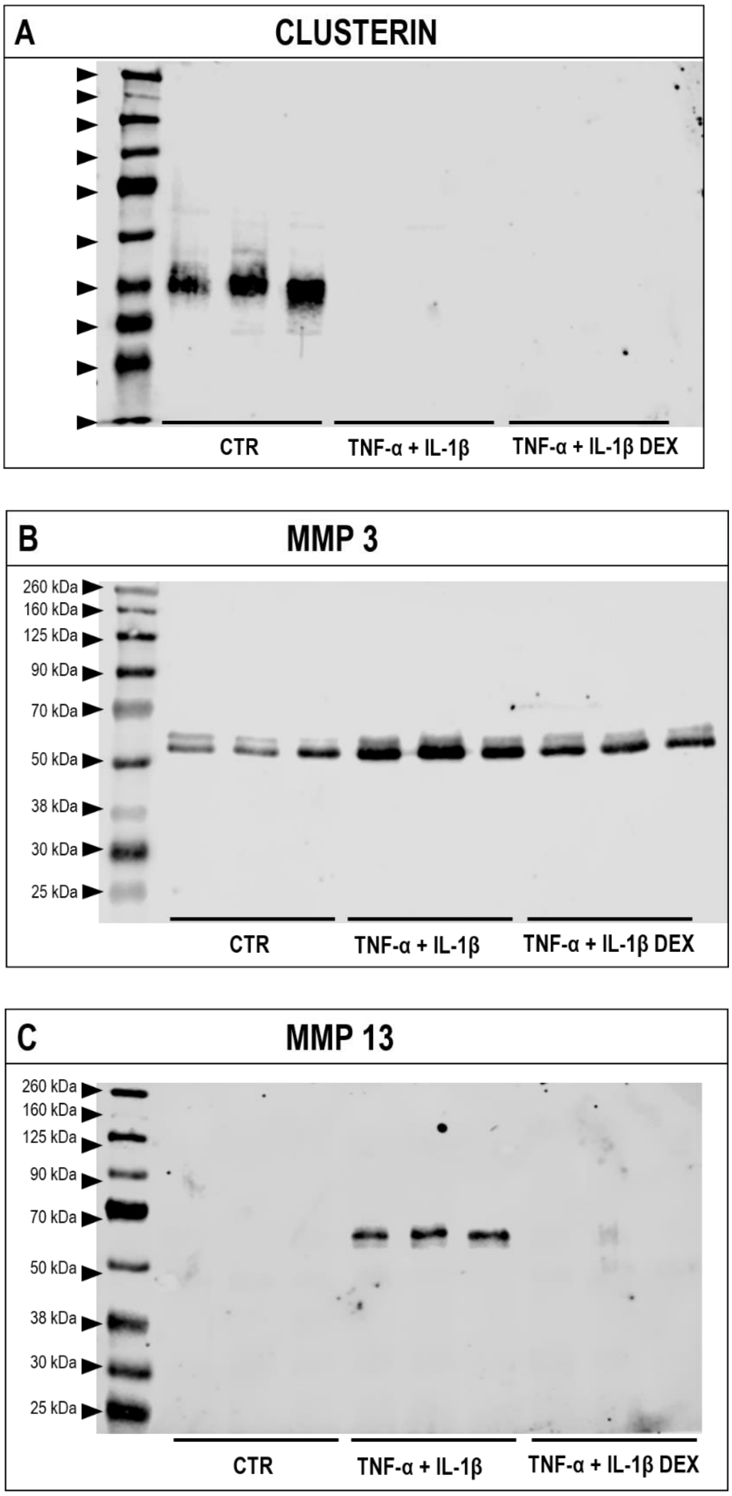


**Figure S4.** Release of **(A)** secreted isoform of clusterin (sCLU), **(B)** matrix metalloproteinase-3 (MMP3) and **(C)** MMP-13 into the secretome of cartilage explants, following stimulation for 7 days with 10 ng/mL TNF-α and 10 ng/mL IL-1β ± 100 μm dexamethasone (DEX). Uncropped membrane images are shown, detected by western blotting and quantified by densitometry.

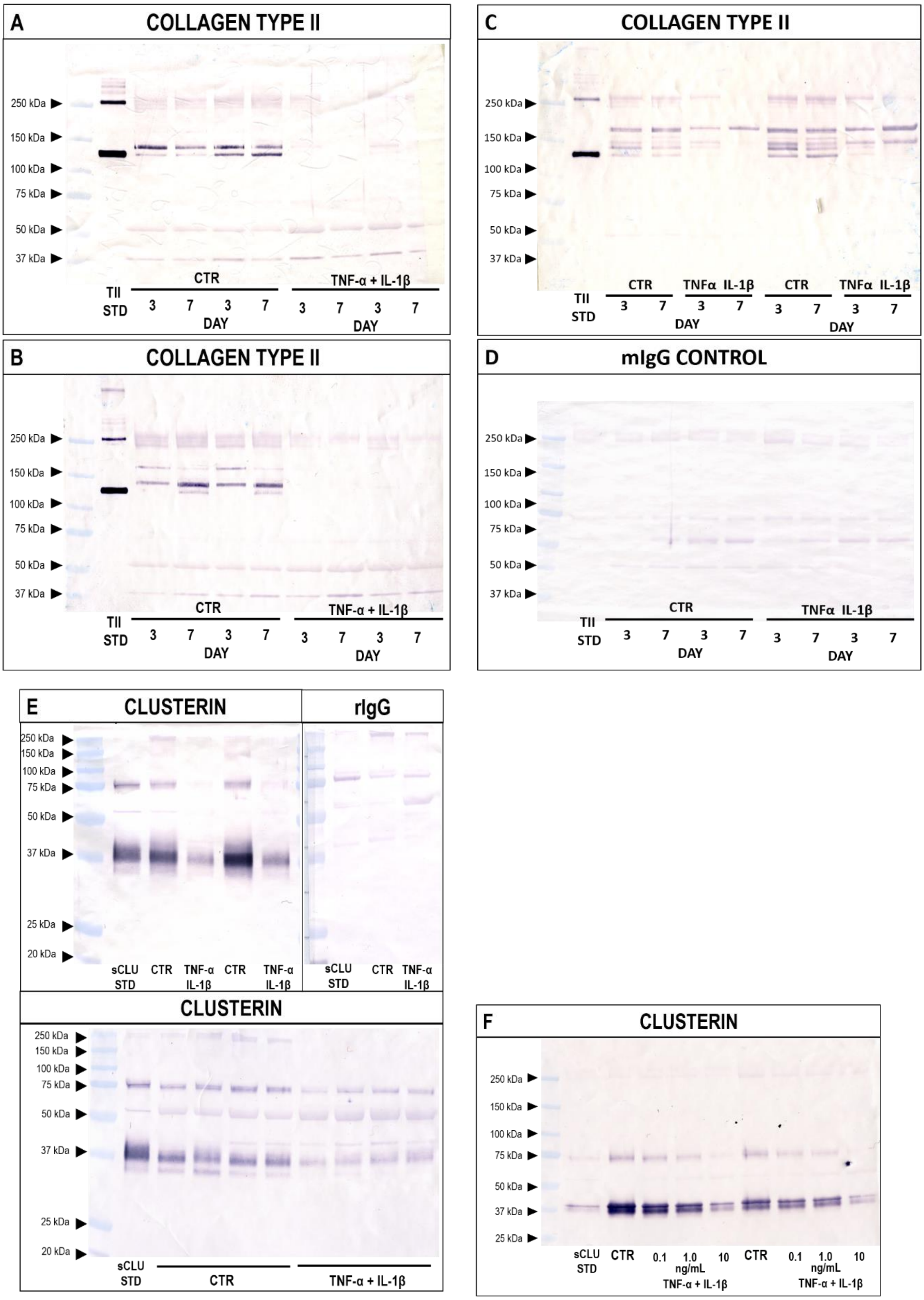


**Figure S5. (A–D)** Release of collagen type II, and **(E–F)** secreted isoform of clusterin (sCLU) into the secretome of un-passaged primary articular chondrocytes, following stimulation with 10 ng/mL TNF-α and 10 ng/mL IL-1β for 7 days. Uncropped membrane images are shown, along with **(D)** mouse (mIgG) and **(E)** rabbit IgG (rIgG) controls, detected by western blotting and quantified by densitometry.

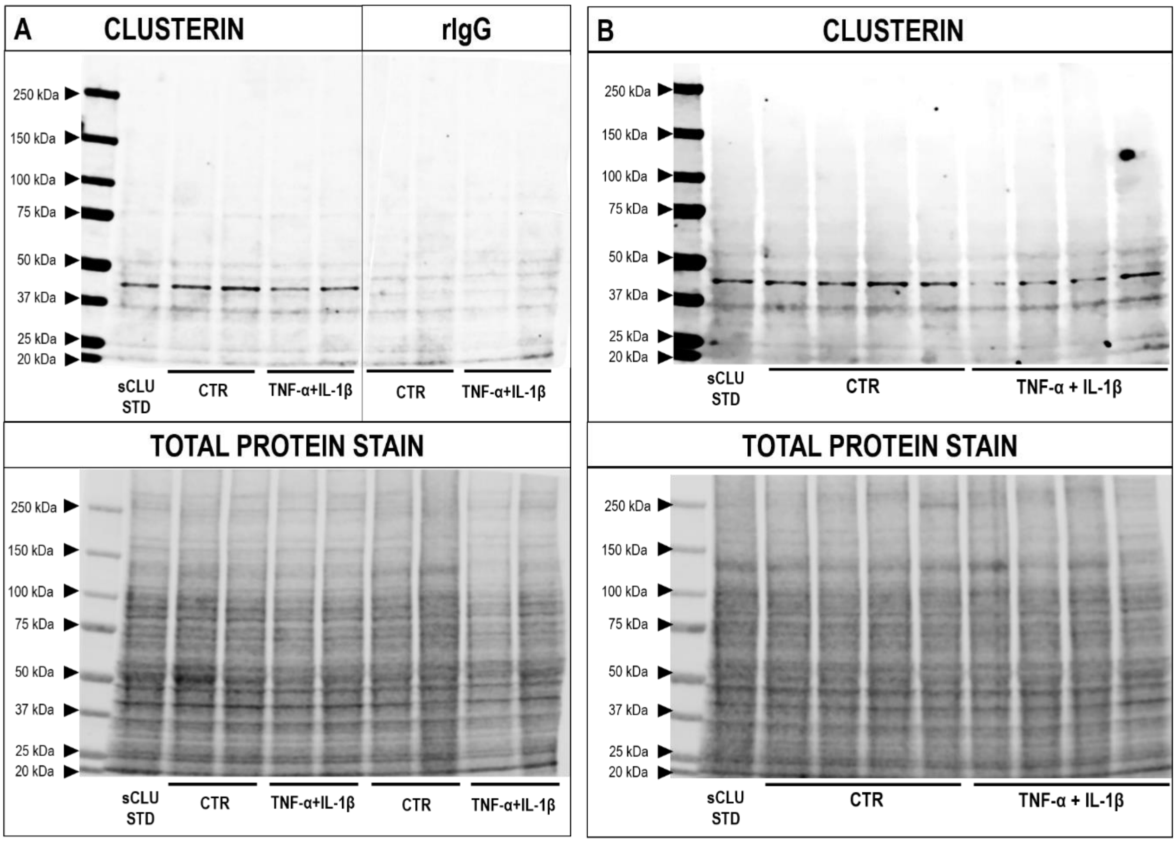


**Figure S6. (A)** Secreted isoform of clusterin (sCLU), and rabbit IgG (rIgG) control, detected by western blotting and quantified by densitometry; normalised to total protein loaded. **(B)** Clusterin protein expression in un-passaged primary articular chondrocytes, following stimulation for 7 days with 10 ng/mL TNF-α and 10 ng/mL IL-1β; normalised to total protein loaded. Uncropped membrane images are shown.

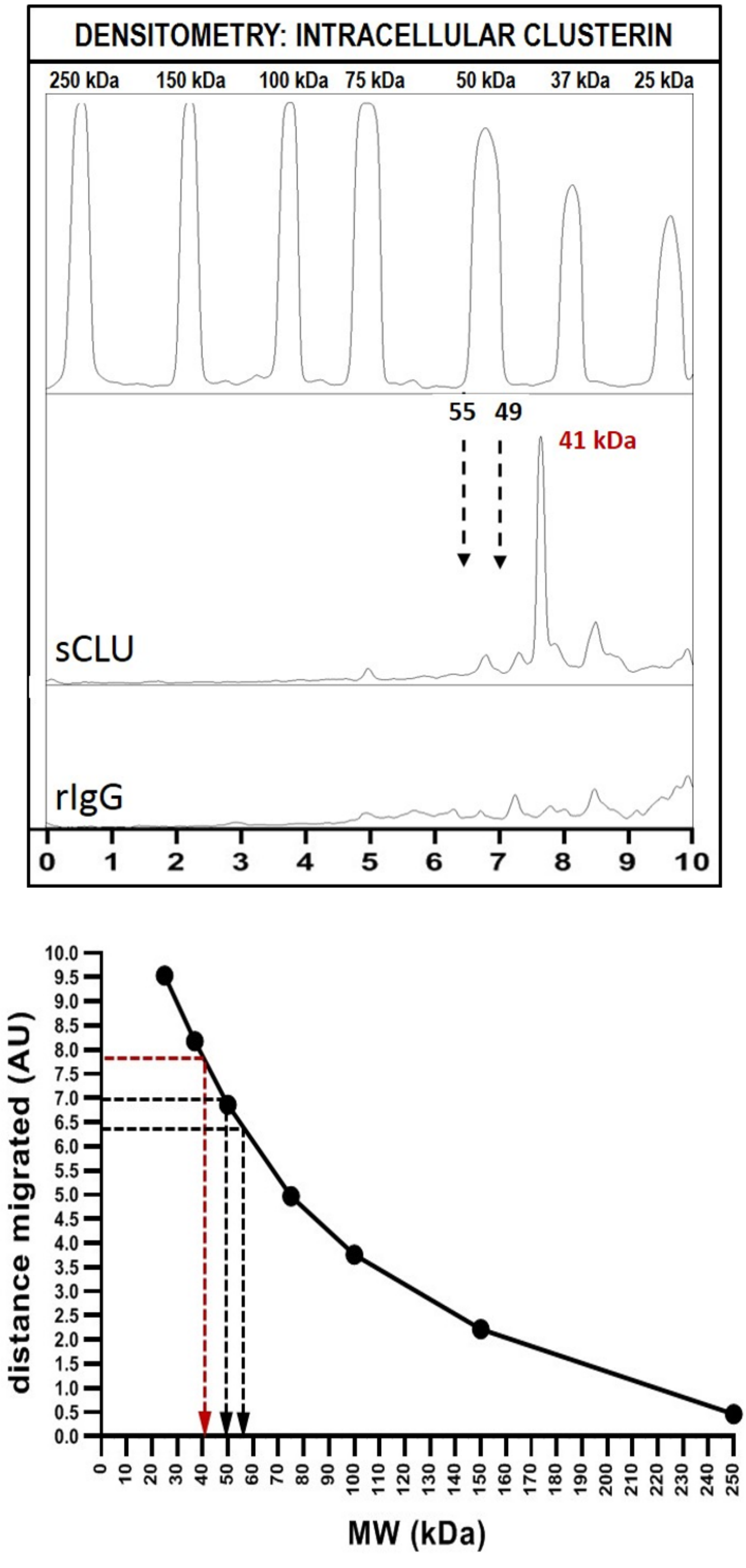


**Figure S7.** Clusterin protein expression in un-passaged primary articular chondrocytes, following stimulation for 7 days with 10 ng/mL TNF-α and 10 ng/mL IL-1β. Secreted isoform of clusterin (sCLU) detected by western blotting and quantified by densitometry. Molecular weight analysis of the band corresponding to clusterin was confirmed at 41.1 kDa (± 0.2), indicating the presence of the intracellular CLU isoform.
